# The Contribution of Brain Structural and Functional Variance in Predicting Age, Sex and Treatment

**DOI:** 10.1101/2020.08.28.272476

**Authors:** Ning-Xuan Chen, Gui Fu, Xiao Chen, Le Li, Michael P. Milham, Su Lui, Chao-Gan Yan

**Affiliations:** CAS Key Laboratory of Behavioral Science, Institute of Psychology, Beijing, China; Magnetic Resonance Imaging Research Center, Institute of Psychology, Chinese Academy of Sciences, Beijing, China; International Big-Data Center for Depression Research, Chinese Academy of Sciences, Beijing, China; Department of Psychology, University of Chinese Academy of Sciences, Beijing, China; Department of Radiology, Sun Yat-sen University Cancer Center, State key laboratory of Oncology in South China, Collaborative Innovation Center for Cancer Medicine; Center for Cognitive Science of Language, Beijing Language and Culture University, Beijing, China; MATTER Lab, Child Mind Institute, New York, NY 10022, USA; Center for Biomedical Imaging and Neuromodulation, Nathan S. Kline Institute for Psychiatric Research, Orangeburg, NY 10962; Department of Radiology, West China Hospital, Sichuan University; Department of Child and Adolescent Psychiatry, Hassenfeld Children’s Hospital at NYU Langone, New York, NY, USA

**Keywords:** model fit, variance partitioning, structural metrics, functional metrics, voxel-based analysis

## Abstract

Structural and functional neuroimaging have been widely used to track and predict demographic and clinical variables, including treatment outcomes. However, it is often difficult to directly establish and compare the respective weights and contributions of brain structure and function in prediction studies. The present study aimed to directly investigate respective roles of brain structural and functional indices, along with their contributions in the prediction of demographic variables (age/sex) and clinical changes of schizophrenia patients. The present study enrolled 492 healthy people from Southwest University Adult Lifespan Dataset (SALD) for demographic variables analysis and 42 patients with schizophrenia from West China Hospital for treatment analysis. We conducted a model fit test with two variables (one voxel-based structural metric and another voxel-based functional metric) and then performed a variance partitioning on the voxels that can be predicted sufficiently. Permutation tests were applied to compare the contribution difference between each pair of structural and functional measurements. We found that voxel-based structural indices had stronger predictive value for age and sex, while voxel-based functional metrics showed stronger predictive value for treatment. Therefore, through variance partitioning, we could clearly and directly explore and compare the voxel-based structural and functional indices on particular variables. In sum, for long-term change variable (age) and constant biological feature (sex), the voxel-based structural metrics would contribute more than voxel-based functional metrics; but for short-term change variable (schizophrenia treatment), the functional metrics could contribute more.

## 1. Introduction

The diverse functional repertoire of the human brain derives from a relatively static anatomy, and the static brain structure affects individual cognition through dynamical functional activities. Brain structures and functions are always coupling with each other, being shaped by genetic and environmental factors together, and reshaping our cognition and behavior (Batista-Garcia-Ramo and Fernandez-Verdecia, 2018; Park and Friston, 2013). As a result, researchers attempting to understand the neural correlates of variations in human behavior across individuals can choose to study either, brain structure or function. This choice remains a difficult question as some argues structural indices are more test-retest reliable but functional metrics are more sensitive to functional disruptions and short-term changes (Shah et al., 2016; Zuo et al., 2013; Zuo et al., 2019). To date, empirical evidence guiding such a selection remains lack.

A few studies have already explored the brain structure-function relationship. Qing and Gong (2016) used resting-state functional magnetic resonance imaging (R-fMRI) on healthy young adults and found a robust positive linear correlation between voxel-based brain volume and amplitude of low frequency fluctuations (ALFF), suggesting a strong association between structure and function (Qing and Gong, 2016). Yang and colleagues (2016) used a data-driven approach, generalized ranking and averaging independent component analysis (gRAICAR), to determine cross-subject co-variance among a few surface-based functional and structural imaging metrics on healthy human brains. Their results revealed a wider scope in searching of multi-modal imaging features, which might be helpful to explain structural substrates of the functional metrics (Yang et al., 2016). Honey and colleagues (2009) used computational modeling to explore the relationship between human resting-state functional connectivity and structural connectivity. They found that although functional connectivity was variable and had few structural linkage, some functional connectivity attributes (strength, persistence, and spatial statistics) were still associated with large-scale cerebral cortex structure (Honey et al., 2009). In addition, other methods, such as generative models (Betzel et al., 2016), network communication theory (Goni et al., 2014), partial least squares (Misic et al., 2016), or simply fitting in a linear/nonlinear stochastic model (Deco et al., 2009; Hansen et al., 2015), had also been used to estimate the function-structure correlation (Wirsich et al., 2017). Although previous researches have clearly implicated a sophisticated relationship between structural and functional features of brain, to what extend they correspond to different human individual characteristics remains largely unexplored.

Some individual characteristics may be particularly intriguing in unraveling different contributions of structural and functional features. First, aging is associated with changes in brain morphology as well as with a decline in cognitive performance (Nakagawa et al., 2013; Persson et al., 2006). Besides, long-term changes on individual structure and function can be used as a biomarker for predicting aging. Second, subtle brain structure differences exist between males and females (Allen et al., 2003), and cognitive abilities also had different representation for each gender, such as better spatial abilities in men and better verbal skills in women. Thus, it would be meaningful to explore the structure-function relationship for clarifying the sex difference, and further to delineate the pathophysiological mechanisms underlying sex differences in neuropsychiatric disorders (Cosgrove et al., 2007). Finally, schizophrenia has been described as a disorder of both, structural and functional connectivity (Skudlarski et al., 2010). In addition, previous studies found that long-term treatment may change schizophrenia brain morphology (Ho et al., 2011) and the short-time treatment mainly change the function rather than structure (Lui et al., 2010). Therefore, through directly comparing the contribution of the brain structural and functional changes to baseline and post-schizophrenia treatment, it might be helpful to locate the drug sensitivity brain areas for further treatment.

With aforementioned three individual characteristics (age, sex and schizophrenia treatment), we aimed to explore the contribution of voxel-based structural metrics and functional metrics to these individual differences, respectively. Gray matter density (GMD) and gray matter volume (GMV) were derived with voxel-based morphometry (VBM) as structural metrics. GMD is the probability of voxels as gray matter versus white matter and other neural tissue, and GMV is the modulated GMD images by using the Jacobian determinants and mirrors volumetric deformation (Ashburner and Friston, 2005). To explore the fundamental properties of intrinsic brain activity, we picked functional metrics including ALFF and fractional ALFF (fALFF), degree centrality (DC), regional homogeneity (ReHo), and voxel-mirrored homotopic connectivity (VMHC) in this study (Yan et al., 2017). On the one hand, as amplitude-based measures had already received widespread attention and could reflect the regional fluctuation characteristics of the intrinsic brain activity, we chose to use ALFF and fALFF to quantify low-frequency characteristics of fluctuations in intrinsic brain activity (Zang et al., 2007; Zuo et al., 2012). On the other hand, we also picked inter-regional synchronization metrics such as DC, ReHo and VMHC, which DC revealed the brain functional organizations in a graph-theoretical way (Buckner et al., 2009; Zuo et al., 2012), ReHo represented the local synchronization of low-frequency fluctuations (Jiang and Zuo, 2016; Zang et al., 2004), and VMHC characterized the functional connectivity between paired symmetric inter-hemispheric voxels (Anderson et al., 2011; Zuo et al., 2010). Although all of these five functional metrics are widely used in the R-fMRI literatures (e.g., see review in Zuo and Xing, 2014), they had always been explored separately and lacked the proof about how similar or distinct between each other(Yan et al., 2017). Therefore, we used all the aforementioned functional measures to provide a comprehensive insight on the interdependencies among them.

The goal of the present study was to directly compare the unique and shared contribution of voxel-level structural and functional metrics in the prediction of a given dependent variable (age, sex or schizophrenia treatment). In each independent prediction model, we used one of structural metrics (GMV and GMD) and one of functional metrics (ALFF, fALFF, DC, ReHo and VMHC) as two independent variables. We utilized a model fit test with both the structural and functional metrics to explore how they could predict on a given variable at voxel-level. Then within the statistically significant brain areas, we perform variance partitioning on each voxel to get four components: the portion of unique variance contributed by structural metrics, the portion of unique variance contributed by functional metrics, the portion of shared variance contributed by the two metrics, and the portion of unexplainable residual variance. Therefore, we could then compare the structural and functional contribution to predict the dependent variable, as well as their shared contribution. Here we hypothesized that for the long-term change variable (age) and the constant biological characteristic variable (sex), the voxel-based structural metrics would contribute more than functional metrics; but for the short-term change variable (schizophrenia treatment), the functional metrics would contribute more.

## 2. Methods

### 2.1. Participants

We performed our analyses on two independent datasets. The first is Southwest University Adult Lifespan Dataset (SALD) which is publicly available. It comprises a large cross-sectional sample (n = 492; 305 Females, 187 Males; age range: 19-80 years) undergoing a multi-modal investigation of these neural underpinnings. All participants were recruited as healthy adult and had no history of psychiatric disorder or use of psychiatric drugs (Wei et al., 2018).The dataset collection was approved by the Research Ethics Committee of the Brain Imaging Center of Southwest University, in accordance with the Declaration of Helsinki. Written informed consent was obtained from all participants prior to the data collection.

The second dataset is from West China Hospital, Sichuan University. This dataset contains 39 patients (22 Females, 17 Males; age range: 16-49 years) with schizophrenia undergoing structural and R-fMRI brain scans before and after a 6-week intervention with antipsychotic medication according to the clinician’s preference, including risperidone, olanzapine, clozapine, quetiapine fumarate, sulpiride and aripiprazole. The study was approved by the Institutional Review Board (IRB) of West China Hospital and all subjects gave written informed consent to their participation. Diagnoses were determined using the Structured Clinical Interview for *DSM-IV* Patient Edition and confirmed after at least 1-year follow-up. All patients were evaluated and scanned at baseline and 6 weeks after treatment.

### 2.2 Imaging protocols

The first dataset was collected at the Southwest University Center for Brain Imaging using a 3T Siemens Trio MRI scanner (Siemens Medical, Erlangen, Germany). Each participant took part in 3D structural MRI (sMRI) and R-fMRI scans. For each participant, the 3D sMRI and R-fMRI sequences were acquired in succession within one session. Three-dimensional T1-weighted magnetization-prepared rapid gradient echo (MPRAGE) sagittal images were obtained using the following sequence: repetition time (TR) = 1,900 ms, echo time (TE) = 2.52 ms, inversion time (TI) = 900 ms, flip angle = 90 degrees, resolution matrix = 256×256, slices = 176, thickness = 1 mm, and voxel size = 1×1×1 mm^3^. Functional images were collected axially using gradient echo echo-planar-imaging (GRE-EPI) sequences: slices = 32, TR = 2000 ms, TE = 30 ms, flip angle = 90 degrees, field of view (FOV) = 220 mm, thickness/slice gap = 3/1 mm, and voxel size = 3.4×3.4×4 mm^3^. Prior to the scan, the subjects were instructed to lie down, close their eyes, and rest without thinking about any specific thing but to refrain from falling asleep. The scan lasted for about 8-min and thus included 242 functional volumes for each subject.

The second dataset was collected from the Department of Radiology, West China Hospital using a 3T GE MR imaging system (EXCITE, General Electric, Milwaukee, USA) with an eight-channel head coil. Participants completed a high-resolution T1-weighted (spoiled gradient sequence, SPGR) sequence and a R-fMRI sequence. The scanning parameters of anatomical images were: TR = 8.528 ms, TE = 3.4 ms, TI = 400 ms, flip angle = 12°, axial slices = 156, FOV = 240 mm, thickness = 1 mm, and voxel size = 0.47×0.47×1 mm^3^. The scanning parameters of R-fMRI were: TR = 2000 ms, TE = 30 ms, flip angle = 90 degrees, voxel size = 3.75×3.75×5 mm^3^, FOV = 240×240 mm. All subjects were instructed to lie down, close their eyes, and keep awake. The scan lasted for 6.5-min and thus included 195 functional volumes for each subject. All images were reviewed by two experienced neuroradiologists to exclude those with gross brain abnormalities and motion artifacts.

### 2.3 Preprocessing

The Data Processing Assistant for Resting-State fMRI (DPARSF, http://rfmri.org/DPARSF) (Yan and Zang, 2010) was used to perform preprocessing. It is based on Statistical Parametric Mapping (SPM12, http://www.fil.ion.ucl.ac.uk/spm) and is integrated in the toolbox for Data Processing & Analysis of Brain Imaging (DPABI, http://rfmri.org/DPABI) (Yan et al., 2016). Firstly, the initial 10 volumes were discarded, then slice-timing correction and head motion realignment were performed. Next, individual T1-weighted MPRAGE images were co-registered to the mean functional image using a 6 degree-of-freedom linear transformation without re-sampling and then segmented into gray matter, white matter (WM), and cerebrospinal fluid (CSF) (Ashburner and Friston, 2005). Finally, transformations from individual native space to MNI space were computed with the Diffeomorphic Anatomical Registration Through Exponentiated Lie algebra tool (Ashburner, 2007).

### 2.4 Nuisance regression

To minimize head motion confounds, we utilized the Friston 24-parameter model (Friston et al., 1996) to regress out head motion effects, which was chosen based on prior work that higher-order models remove head motion effects better (Satterthwaite et al., 2013; Yan et al., 2013). Additionally, mean FD was used to address the residual effects of motion in group analyses. Mean FD is derived from Jenkinson’s relative root mean square algorithm (Jenkinson et al., 2002). WM and CSF signals (using DPARSF’s default WM and CSF masks) were also removed from the data through linear regression. Additionally, linear trends were included as a regressor to account for drifts in the blood oxygen level dependent signal. We performed temporal bandpass filtering (0.01–0.1 Hz) on all time series except for ALFF and fALFF analyses.

### 2.5 Structural metrics

We used two voxel-based structural metrics including GMV maps and GMD maps to do further analysis. GMV was modulated GMD images by using the Jacobian determinants derived from the spatial normalization in the voxel-based morphometry (VBM) analysis (Good et al., 2001). The GMV and GMD maps were smoothed by a full width at half maximum (FWHM) of 4mm. In order to perform the structural-functional contribution analysis, we resliced the structural metrics to the resolution of functional metrics, i.e., 3×3×3 mm^3^.

### 2.6 Functional metrics

Here we used five kinds of voxel-based functional metrics (ALFF, fALFF, DC, ReHo and VMHC) to represent different functional aspects.

ALFF is the mean amplitude of low-frequency fluctuations (0.01–0.1Hz) by the fast Fourier transform in the time course of each voxel (Zang et al., 2007).

fALFF is a normalized version of ALFF and calculated as the total power in the low-frequency range (0.01–0.1Hz) divided by the whole power of entire frequency range of the same voxel (Zou et al., 2008).

DC is the number or sum of significant connections’ weights for each voxel. The weighted sum of positive correlations was calculated, including only those connection’s correlation coefficient exceeding the threshold of *r* = 0.25 was kept (Buckner et al., 2009; Zuo et al., 2012).

ReHo is the homogeneity of a voxel’s timeseries with its nearest local neighbors’ (26 voxels used here) time courses. It is calculated as Kendall’s coefficient of concordance (KCC) (Zang et al., 2004).

VMHC indicates the functional connectivity between paired symmetric inter-hemispheric voxels. It is calculated as the Pearson’s correlation coefficient between the time series of one voxel and that of its counterpart voxel located the same in the opposite hemisphere. Then, the values of VMHC were Fisher’s *r*-to-*z* transformed. For better correspondence for counterpart voxels, individual functional data were further registered to a group averaged symmetric template, which was created by computing a mean normalized T1 image across all the participants, and then averaged with correspondent left–right mirrored version (Anderson et al., 2011; Zuo et al., 2010).

All these metrics are Z-standardized (i.e., subtract the whole brain mean and then divided by the whole brain standard deviation) for each subject. Then we smoothed (4 mm FWHM) all the R-fMRI Metrics, except for VMHC as they were smoothed and Fisher-Z transformed beforehand.

### 2.7 Statistical analysis for model fit

We used one of the two structural parameters including GMV and GMD and one of the five functional metrics including ALFF, fALFF, DC, ReHo and VMHC as two independent variables (predictors) (Chen et al., 2018; Yan et al., 2017). We then used these two independent variables to predict age or sex in the first dataset and treatment timepoint (baseline or post-treatment) in the second dataset. Specifically, for our study, the following regression model was constructed:

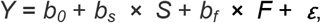

where *Y* is one of the three dependent variables (age, sex or treatment), *b_0_* is the intercept parameter, *S* denotes one of two structural parameters (GMV or GMD), *b_s_* is the regression coefficient corresponding to *S*, *F* denotes one of the five functional metrics (ALFF, fALFF, DC, ReHo or VMHC), *b_f_* is the regression coefficient corresponding to *F*, and *ε* is the residual. Therefore, we separately performed an F-test for the model fit with each model so that ten F maps were generated for each of the three dependent variables. Then one-tailed gaussian random field (GRF) correction was used for multiple comparison correction on these F maps with voxel-level thresholds of *p* < 0.001 and cluster-level thresholds of *p* < 0.05, as recent studies suggest strict voxel-level p threshold (cluster defining threshold) should be adopted (Chen et al., 2018; Eklund et al., 2016).

### 2.8 Variance partitioning

To further identify the unique and shared variances of the structural and functional indices for predicting a given dependent variable, a voxel-level variance partitioning was conducted on the voxels those showed a significant model fit. We performed the variance partitioning using the package vegan (version 2.5, Oksanen et al., 2018) in R (version 3.6.0, R Foundation for Statistical Computing, Vienna, Austria). Variance partitioning gave four components of results: the portion of unique variance contributed by structural parameters (V_s_), the portion of unique variance contributed by functional metrics (V_f_), the portion of shared variance contributed between the structural and functional metrics (V_sh_), and the portion of unexplainable residual variance (V_r_). The sum of the four portions is 100%. The portion difference between the unique variances contributed by structural and functional metrics (V_s-f_) was calculated as (V_s_ - V_f_) on each voxel.

### 2.9 Permutation test

For each component of variance partitioning, we used permutation test to infer the statistical significance. To reduce the computational load, we created permutation-based null distributions for 200 regions following the Craddock’s functional clustering atlas (Craddock et al., 2012), and compared each voxel with the null distribution of the region that contains that voxel. For each atlas parcel, we performed 10000 times variance partitioning while exchanging the structural and functional metrics for all the subjects randomly. By comparing each voxel-level variance partitioning value with the null distribution of its related region, we were able to calculate the *p* value and the corresponding *z* value. Finally, we performed one-tailed GRF correction for V_s_, V_f_, and V_sh_, with voxel-level threshold of *p* < 0.001 and cluster-level threshold of *p* < 0.05. The same calculation also applied to V_s-f_, the portion difference between the unique variances contributed by structural and functional metrics.

## 3. Results

### 3.1 Structural and Functional Metrics in Predicting Age

Based on the F-test of model fit, we found that age could be predicted by combining structural and functional measurements for a broad range of brain areas, including precentral gyrus, superior temporal gyrus, insula, thalamus (see Fig. S1). Based on whole brain variance partitioning maps of age, structural metrics contributed significant portion of unique variance on a broad range of brain areas, while neither the functional metrics nor the shared portion of both metrics showed any significant portion of unique variance across the brain. As demonstrated in Fig. 1, the superior temporal gyrus, precentral gyrus, inferior frontal gyrus, middle frontal gyrus, and cingulate gyrus all showed significant V_s_ when using all the two structural metrics with all the five functional metrics to predict age. Furthermore, medial frontal gyrus and superior frontal gyrus also exhibited significant V_s_ only when using GMV and all the five functional metrics to predict age.

**Fig. 1.**
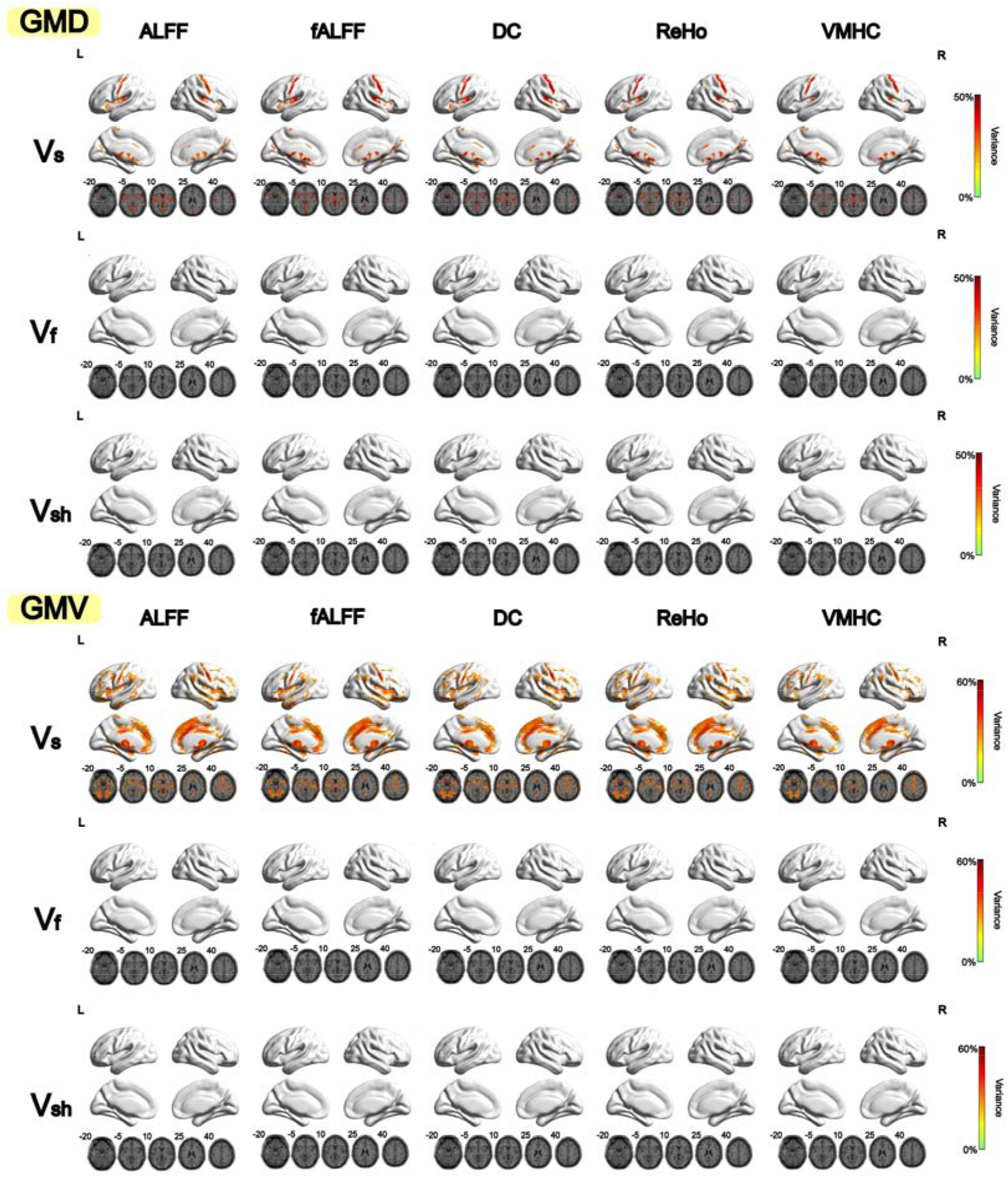
Variance partitioning of voxel-based structural metrics (GMD or GMV) and functional metrics (ALFF, fALFF, DC, ReHo or VMHC) in predicting age (percentages explained of the total variance). For each component of permutation test. GMD stands for gray matter density, and GMV stands for gray matter volume. ALFF stands for amplitude of low frequency fluctuations, fALFF stands for fractional ALFF, DC stands for degree centrality, ReHo stands for regional homogeneity, and VMHC stands for voxel-mirrored homotopic connectivity. V_s_ indicates the portion of unique variance contributed by structural parameters, V_f_ indicates the portion of unique variance contributed by functional metrics, and V_sh_ indicates the portion of shared variance contributed between the structural and functional metrics.

For age, based on portion difference between the unique variances contributed by structural and functional metrics (see Fig. 2), For most brain regions, V_s_ was significantly higher than V_f_ (significantly positive V_s-f_ values), especially for GMV. We found that superior temporal gyrus, precentral gyrus, inferior frontal gyrus, middle frontal gyrus, cingulate gyrus showed significantly more positive V_s-f_ when using either of the two structural metrics and any of the five functional metrics to predict age. Furthermore, medial frontal gyrus and superior frontal gyrus also exhibited significantly more positive V_s-f_ when using GMV and any of the five functional metrics to predict age.

**Fig. 2.**
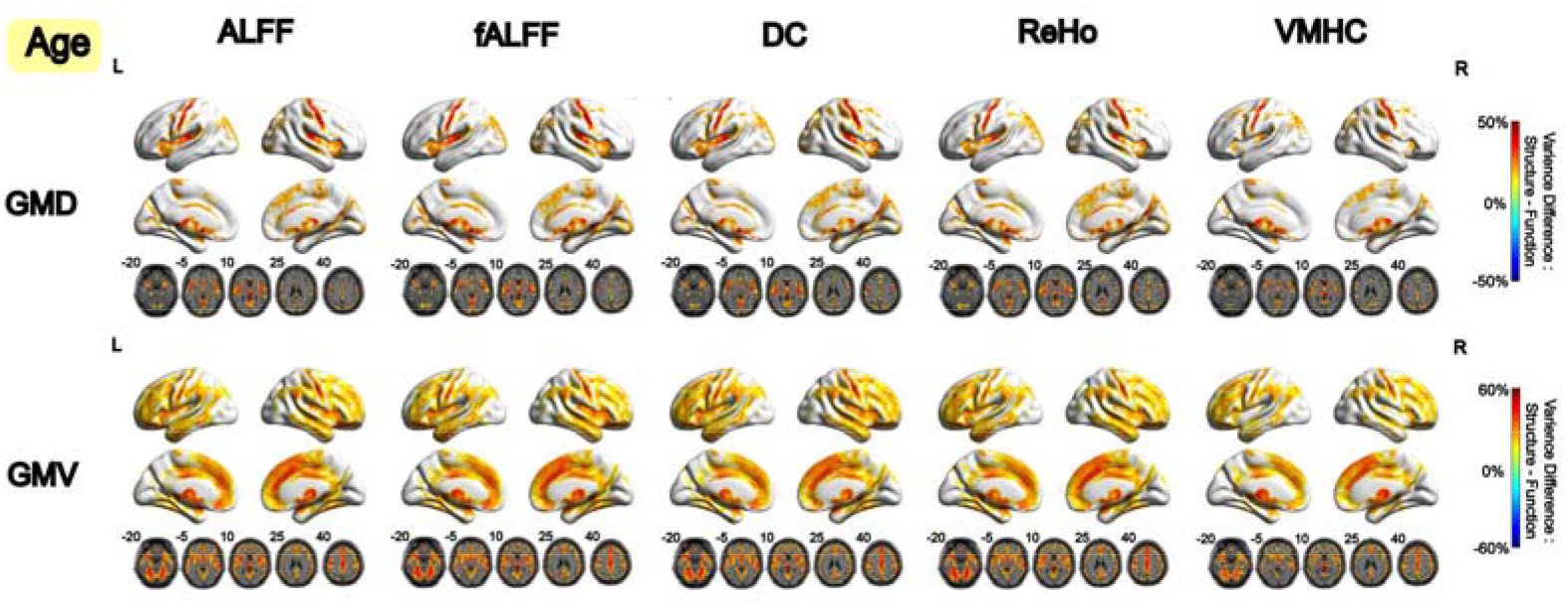
Difference between the unique variances contributed by voxel-based structural metrics (GMD or GMV) and functional metrics (ALFF, fALFF, DC, ReHo or VMHC) (V_s-f_) in predicting age (percentages explained of the total variance). It was calculated as (V_s_ - V_f_) on each voxel. GRF correction (voxel-level *p* < 0.001, cluster-level *p* < 0.05) was utilized to correct the large number of voxels across the brain, while the voxel-level *p* values were calculated by comparing each voxel-level variance partitioning value with the null distribution of its related region generated by permutation test.

### 3.2 Structural and Functional Metrics in Predicting Sex

Based on the F-test of model fit, we found that sex could be predicted by combining structural and functional measurements for a few brain areas: sub-lobar, thalamus, and caudate for GMD and any of the five functional metrics, and sub-lobar, parahippocampal gyrus for GMV and any of the five functional metrics (see Fig. S2). Based on the whole brain variance partitioning maps of sex, structural metrics contributed significant portion of unique variance on a few of brain areas, while the functional metrics and the shared portion of both metrics showed few significant portion of unique variance across the brain. It was observed that thalamus and caudate showed significant V_s_ when using GMD and any of the five functional metrics to predict sex, while cerebellum, occipital lobe, temporal lobe and insula showed significant V_s_ when using GMV and any of the five functional metrics to predict sex (see Fig. 3). An exception is that precentral gyrus showed significant V_f_ when using GMD and ALFF to predict sex.

**Fig. 3.**
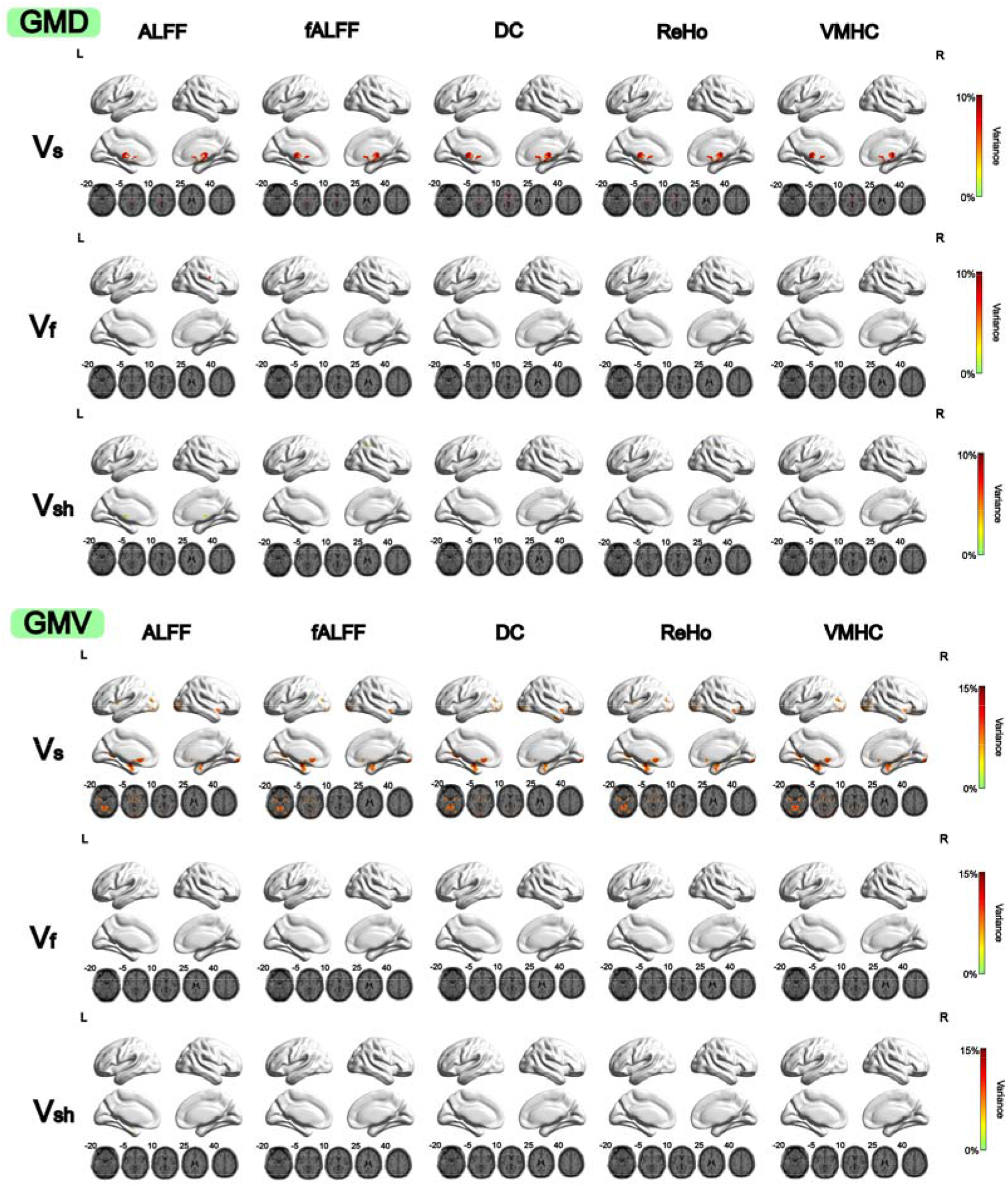
Variance partitioning of voxel-based structural metrics (GMD or GMV) and functional metrics (ALFF, fALFF, DC, ReHo or VMHC) in predicting sex (percentages explained of the total variance). For each component of variance partitioning on predicting sex, GRF correction (voxel-level *p* < 0.001, cluster-level *p* < 0.05) was utilized to correct the large number of voxels across the brain, while the voxel-level *p* values were calculated by comparing each voxel-level variance partitioning value with the null distribution of its related region generated by permutation test.

Furthermore, for sex, based on portion difference between the unique variances contributed by structural and functional metrics (see Fig. 4), V_s_ was also significantly higher than V_f_ across the brain (significantly positive V_s-f_ values). It was observed that thalamus and caudate showed significantly more positive V_s-f_ when using GMD and any of the five functional metrics to predict sex, while cerebellum, occipital lobe, temporal lobe and insula showed significantly more positive V_s-f_ when using GMV and any of the five functional metrics to predict sex. An exception is that precentral gyrus showed more negative V_s-f_ when using GMD and ALFF to predict sex.

**Fig. 4.**
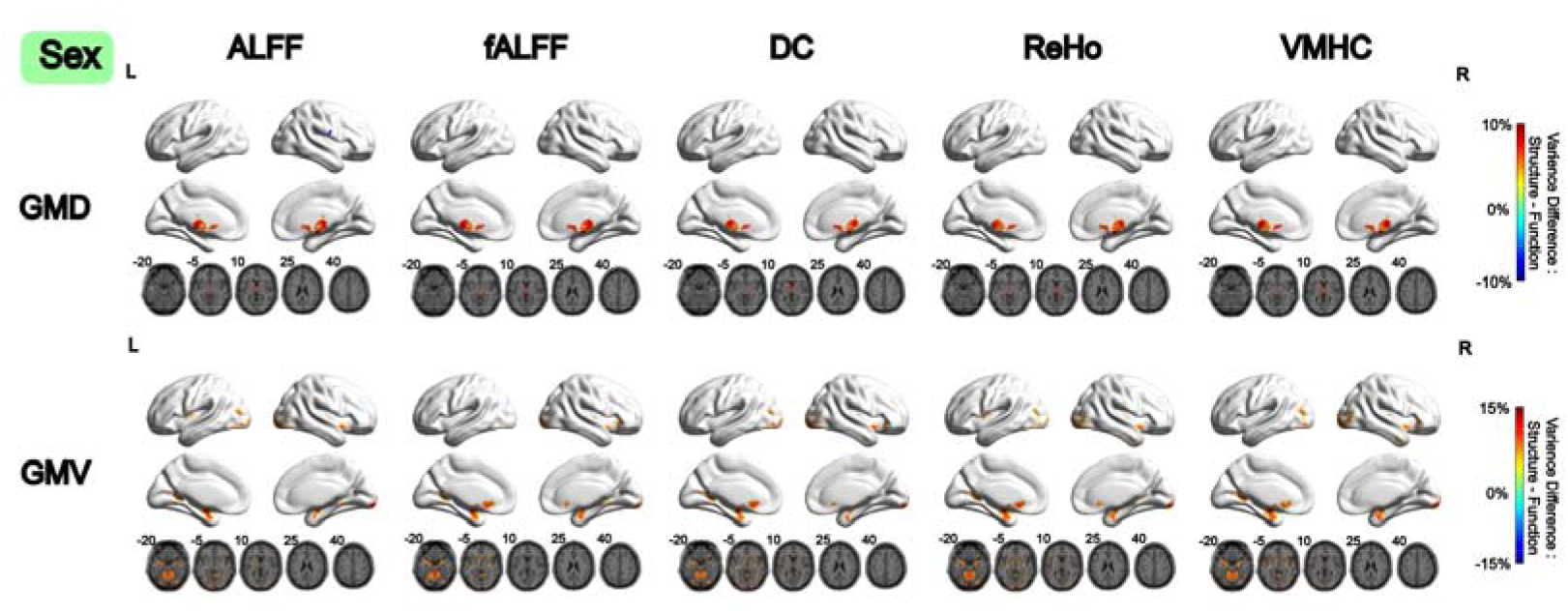
Difference between the unique variances contributed by voxel-based structural metrics (GMD or GMV) and functional metrics (ALFF, fALFF, DC, ReHo or VMHC) (V_s-f_) in predicting sex (percentages explained of the total variance). It was calculated as (V_s_ - V_f_) on each voxel. GRF correction (voxel-level *p* < 0.001, cluster-level *p* < 0.05) was utilized to correct the large number of voxels across the brain, while the voxel-level *p* values were calculated by comparing each voxel-level variance partitioning value with the null distribution of its related region generated by permutation test.

### 3.3 Structural and Functional Metrics in Predicting Treatment (pre- or post-treatment)

Based on the F-test of model fit, we found that pre- and post-treatment could be predicted by combining structural and functional measurements for a few brain areas, which frontal lobe showed a prediction effect by using GMD with fALFF or VMHC, and cerebellum and frontal lobe showed a prediction effect by using GMV with fALFF or VMHC (see Fig. S3). According to whole brain variance partitioning maps of treatment, functional metrics contributed significant portion of unique variance on a few brain areas, while neither the structural metrics nor the shared portion of both metrics showed any significant portion of unique variance across the brain. It was observed that occipital lobe, lingual gyrus showed significant V_f_ when using all the two structural with fALFF or VMHC to predict treatment, while right cerebellum showed significant V_f_ when using GMV with fALFF or VMHC to predict treatment (Fig. 5).

**Fig. 5.**
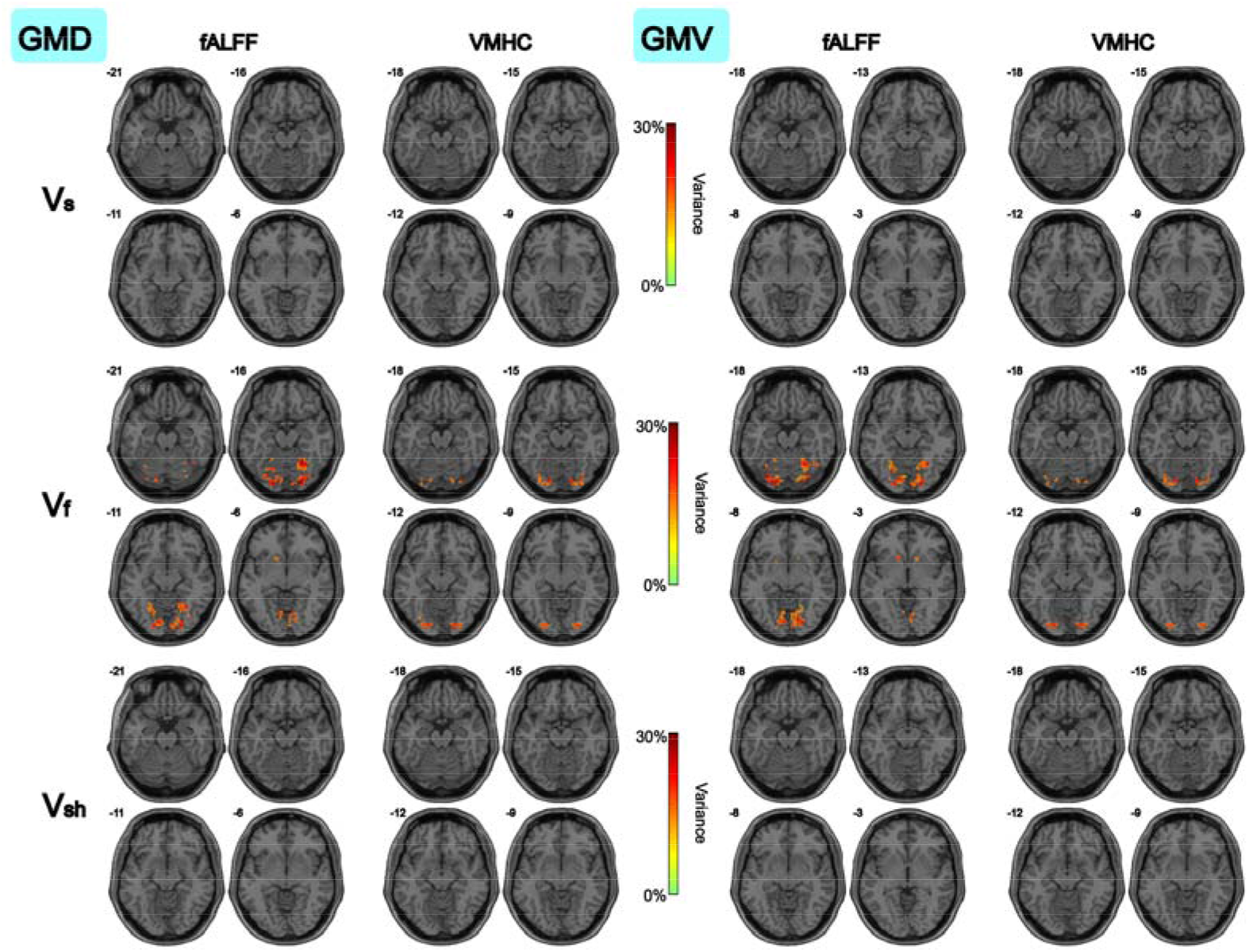
Variance partitioning of voxel-based structural metrics (GMD or GMV) and functional metrics (fALFF or VMHC) in predicting treatment (percentages explained of the total variance). For each component of variance partitioning on predicting treatment time, GRF correction (voxel-level *p* < 0.001, cluster-level *p* < 0.05) was utilized to correct the large number of voxels across the brain, while the voxel-level *p* values were calculated by comparing each voxel-level variance partitioning value with the null distribution of its related region generated by permutation test.

However, for treatment, based on portion difference between the unique variances contributed by structural and functional metrics (see Fig. 6), V_f_ was significantly higher than V_s_ across the brain (significantly negative V_s-f_ values). It was observed that occipital lobe, lingual gyrus showed significantly more negative V_s-f_ when using all the two structural metrics with fALFF or VMHC to predict treatment.

**Fig. 6.**
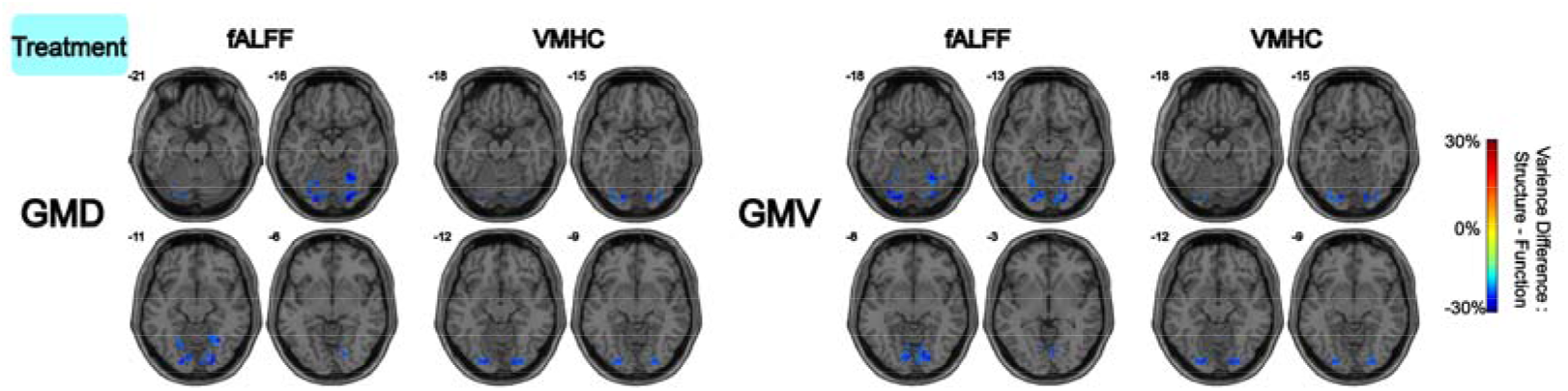
Difference between the unique variances contributed by voxel-based structural metrics (GMD or GMV) and functional metrics (ALFF or VMHC) (V_s-f_) in predicting treatment (percentages explained of the total variance). It was calculated as (V_s_ - V_f_) on each voxel. GRF correction (voxel-level *p* < 0.001, cluster-level *p* < 0.05) was utilized to correct the large number of voxels across the brain, while the voxel-level *p* values were calculated by comparing each voxel-level variance partitioning value with the null distribution of its related region generated by permutation test.

## 4. Discussion

As most studies focused on separately exploring the structural and functional metrics on certain variables, it would be intriguing to combine both to achieve a more thorough explanation of brain change. In our study, we directly tested two sets of independent voxel-based variables (structural vs. functional) in their model fit on age, sex and treatment. Variance partitioning was also performed to compare the portions of explained variances contributed by the structural and functional measurements. We found that on age and sex, structural indices had a stronger predicting effect than functional metrics; but on treatment, the functional metrics showed a stronger predicting effect than the structural ones. Our results indicated that age and sex differences may be better characterized by structural features, while relatively short-term alterations like treatment effects may be better revealed by functional metrics, while all metrics were voxel-based derived in volume space. Therefore, through variance partitioning, we could clearly and directly explore and compare the structural and functional indices on particular variables. The choice of using structural or the functional metrics should depend on the research topics.

### 4.1 Comparing the contributions of structural and functional metrics in a comprehensive model

First of all, the goal of testing the model fit was to combine the structural and functional effect together to attain a better explanation than simply using a single one. In this way, we not only get the unique effect of the voxel-based structural and functional indexes contributing on significant brain areas, but also the shared component. Thus, the difference between these two sets of brain features can be evaluated and compared in a more comprehensive way and let us determine the common and unique contributions from them on predicting a given variable. Moreover, such a two-variable model fit test can model more effects on certain variables than a traditional single-variable model.

Variance partitioning was also performed on brain areas whose metrics can significantly predict our tested “dependent variables” (age, sex and treatment). Variance partitioning has been proved to be a powerful avenue to compare different sources of variance for a certain variable (Groen et al., 2018; Lescroart et al., 2015). For example, Zhao and colleagues (2019) used Joint and Individual Variation Explained (JIVE) method to explore what extant different cortical shape measures covary. They also found the covariation pattern had an excellent prediction effect on age, sex and IQ (Zhao et al., 2019). However, this method only focused on the shared covary component but neglect the unique components, nor take predicting a target variable into account in estimating shared covariation. In our study, we compared both the unique components and the shared component. We found that, for age and sex, V_s_ was higher than V_f_ and V_sh_; but for schizophrenia treatment, V_f_ was higher than V_s_ and V_sh_. Therefore, with the method, we could separately inspect the predicting effect of each component from variance partitioning, and it can further give us advice about how to select structural or functional metrics on different research topics.

Importantly, a new index as the contribution difference between the voxel-based structural and functional metrics was calculated in our study, namely V_s-f_, by which V_s_ subtracted V_f_. This index offered a direct way to compare the structural and functional effect on given variables. It revealed that for age and sex, the V_s-f_ was more positive which indicated the structural indices had more contribution than the functional ones; while for treatment outcome, the V_s-f_ was more negative which meant the functional indices contributed more. Therefore, with this new index V_s-f_, comparing the contribution between voxel-based structural and functional measurements came to be in a unified way which could provide new perspectives.

### 4.2 Selecting voxel-based structural or functional metrics on different research topics

As aging leads to widespread neurobiological changes, it would impact the structural organization and integrity on which large-scale networks critically depend (Greicius et al., 2009; Horn et al., 2014; Teipel et al., 2010; van den Heuvel et al., 2008). Thus, structural changes related to aging would, in part, account for the functional changes associated with cognitive decline. In addition, it has been suggested that older adults’ brains compensate for a reduced structural integrity through increased functional activity (Grady et al., 2010; Park and McDonough, 2013). However, from our observations on voxel-based metrics derived in volume space, brain structure changes accounts significantly more variance than functional ones, thus imply the structural characteristics may develop substantially over age while brain functions remain stable though compensation.

Furthermore, sex differences on brain structure might be related to the differences in sex-specific hormones, but it might also cause the compensatory mechanisms to reduce the functional discrepancy between males and females (De Vries, 2004; McCarthy and Arnold, 2011; Ritchie et al., 2018). This perspective may explain our results that V_s_ was much higher over V_f_. Structural differences reflect sexual biological constant characteristic which indicate more fundamental genetic expression, whereas the unmatched functional results may indicate the widespread functional compensatory through the whole brain between different sex, for example, men and women achieve a similar intelligence quotient (IQ) by the exploitation of different brain areas (Cosgrove et al., 2007). We found that thalamus and caudate showed a higher V_s_ when using GMD and any of the five functional metrics in predicting sex. Men have a larger thalamus, while women have a larger caudate (Cosgrove et al., 2007). The differences between these regions may reveal the different distribution of androgen and estrogen receptors (Cosgrove et al., 2007). Thus, based on voxel-based analysis, the sexual difference has a stronger representation on structure rather than the function.

For the treatment, as all patients with schizophrenia only accepted a 6-weeks treatment in our study, the drugs’ effect would first manifest through functional alterations rather than brain structural changes (Lui et al., 2010). Occipital lobe, lingual gyrus showed higher V_f_ on GMD/GMV with fALFF/VMHC. The occipital lobe is popularly known to be associated with the sense of vision. It may indicate the symptoms such as delusions, hallucinations have been relieved after treatment (Tohid et al., 2015). The higher functional changes on cerebellum of GMV may reflect deficits in motor control seen in patients with schizophrenia (Hoptman et al., 2010). Therefore, for those short-term treatment-related neuroimaging studies, more attention should be paid on the functional changes rather the brain morphology.

### 4.3 Recommendations

Based on the present voxel-based analysis in volume space, some recommendations regarding suitable neuroimaging metrics for different research topics may be given. If variables are long time change or constant biological characteristics, like age, sex or handedness (Fjell et al., 2014; Kertesz et al., 1990), or long-term training (Draganski et al., 2004; Lazar et al., 2005), then voxel-based structural metrics should be recommended; On the other hand, for investigating short-term effects, such as short-term psychiatric treatment outcomes (Lui et al., 2010; Wang et al., 2014), or short-time practice (Erickson et al., 2006; Tomasi et al., 2004), voxel-based functional metrics may be better. It might offer a suggestion about which voxel-based metric could be more effective to explore a specific phenomenon so that it could save time and be more precise.

However, the shared contribution of voxel-based structural and functional metrics generally was small on brain areas with significant model-fit among the three dependent variables used here. This may imply the independent effects of structural and functional metrics are larger than the shared one on age, sex and treatment outcomes. Nevertheless, combining two kinds of metrics can achieve better effects than using an independent variable alone (structural or functional metrics). Although the shared contribution was small on the three dependent variables, the variance-partitioning method still offers a new sight into directly exploring the relationship between voxel-based structural and functional metrics.

In addition, the structural and functional measurements have different reliability levels: the reliability of structural metrics were generally higher than functional ones (Zuo et al., 2019). However, the current study suggests that the structural and functional metrics may have different validity depending on the research questions. Based on our voxel-based analysis, for age and sex, a higher V_s_ may indicate that the two demographic features have been more sensitive to brain demographic feature change; while for schizophrenia treatment, a higher V_f_ indicated that the functional change may be more sensitive than the structural metrics. Thus, beyond considering reliability in selecting a give measure in a specific research topic, attention should be paid to the validity, although which is difficult to be examined. Our index V_s-f_ provides a hint to guide making a choice.

### 4.4 Limitations

Some limitations should be noted for this study. First, here we only performed voxel-based analysis in volume space. It’s suggested that surface-based analyses is spatially more accurate than volume-based analyses (Coalson et al., 2018). We don’t know if our current conclusion can be generalized to vertex-based metrics in surface space, which needs future work. Second, with the variance partitioning, large amount of residual remained, especially for sex and treatment outcomes. It means that only by the MRI indices used here, we cannot fully explain the change of these variables. Therefore, if we consider combining other neuronal measurements (e.g. electroencephalogram or magnetoencephalography), we may obtain a better explanation of variances. Third, here we used the linear model to explore the model fit between the structural and functional metrics, but we cannot detect the nonlinear prediction effect between the variables. Further studies should also consider to model the non-linear effects. Finally, for age, only cross-sectional samples were involved here. Thus, we were not able to examine the longitudinal variation within the participants which might offer more biological meaningful results. Further studies could test the model in longitudinal data.

## Conclusions

In summary, we used variance partitioning to directly compare the structural and functional contribution on age, sex and treatment. The voxel-based structural metrics contributed more on age and sex, while the voxel-based functional metrics had more contribution on schizophrenia treatment. Most importantly, our study offers a new sight that based on voxel-level analysis, for the long-term change variable (age) and constant biological characteristic variable (sex), more attention should be paid to the structural metrics; but for short-term change variable (schizophrenia treatment), we should focus more on the functional measurements.

## Declarations

### Funding

This work was supported by the National Key R&D Program of China (2017YFC1309902), the National Natural Science Foundation of China (81671774, 81630031, 81671664, and 81621003), Beijing Nova Program of Science and Technology (Z191100001119104), the 13th Five-year Informatization Plan (XXH13505) of Chinese Academy of Sciences, the Key Research Program of the Chinese Academy of Sciences (ZDBS-SSW-JSC006), and the 1.3.5 Project for Disciplines of Excellence, West China Hospital, Sichuan University (Project No. ZYYC08001, ZYJC18020). Dr. Su Lui also acknowledge the support from Humboldt Foundation Friedrich Wihelm Bessel Research Award.

### Conflicts of interest/Competing interests

The authors declare no competing financial interests.

### Ethics approval and Consent to participate

The first dataset of Southwest University Adult Lifespan Dataset (SALD) is publicly available. All data collected was approved by the Research Ethics Committee of the Brain Imaging Center of Southwest University and only after informed consent was obtained. The second study about patients with schizophrenia from West China Hospital, Sichuan University was performed with approval of the Institutional Review Board (IRB) of West China Hospital and all subjects gave written informed consent to their participation.

### Consent for publication

Written informed consent for publication was obtained from all participants.

### Availability of data and material

The first dataset of Southwest University Adult Lifespan Dataset (SALD) is publicly available from the International Data-sharing Initiative (http://fcon_1000.projects.nitrc.org/indi/retro/sald.html). The second dataset of patients with schizophrenia from West China Hospital, Sichuan University is not available in the public domain due to patient privacy concerns.

### Code availability

The Data Processing Assistant for Resting-State fMRI (DPARSF) was used to perform preprocessing and the source code of it is openly shared at http://rfmri.org/DPARSF.

### Authors’ contributions

Conception and design: Ning-Xuan Chen, Michael P. Milham and Chao-Gan Yan. Data collection: Gui Fu and Su Lui. Data analysis: Ning-Xuan Chen, Le Li, and Chao-Gan Yan. Drafting of the manuscript: Ning-Xuan Chen, Gui Fu, Xiao Chen, Michael P. Milham, Su Lui, Chao-Gan Yan.

## Notes

### Competing Interest Statement

The authors have declared no competing interest.

