## Supplementary material for "The Contribution of Brain Structural and Functional Variance in Predicting Age, Sex and Treatment": Fig. S1-3

### Supplement Material

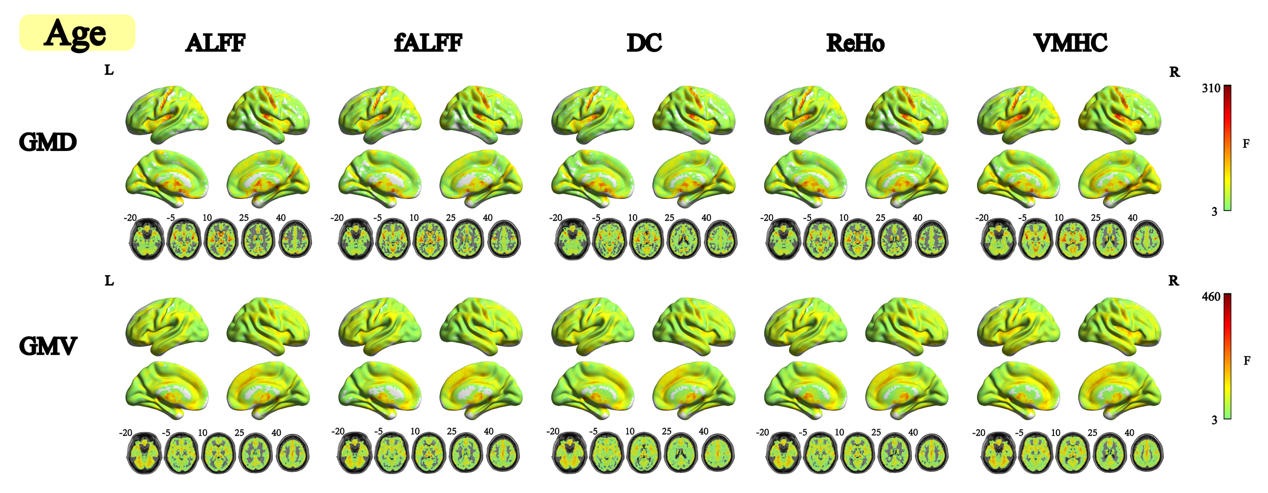

Fig. S1 F-test results of age. GRF correction (voxel-level *p* < 0.001, cluster-level *p* < 0.05) was utilized to correct the large number of voxels across the brain.

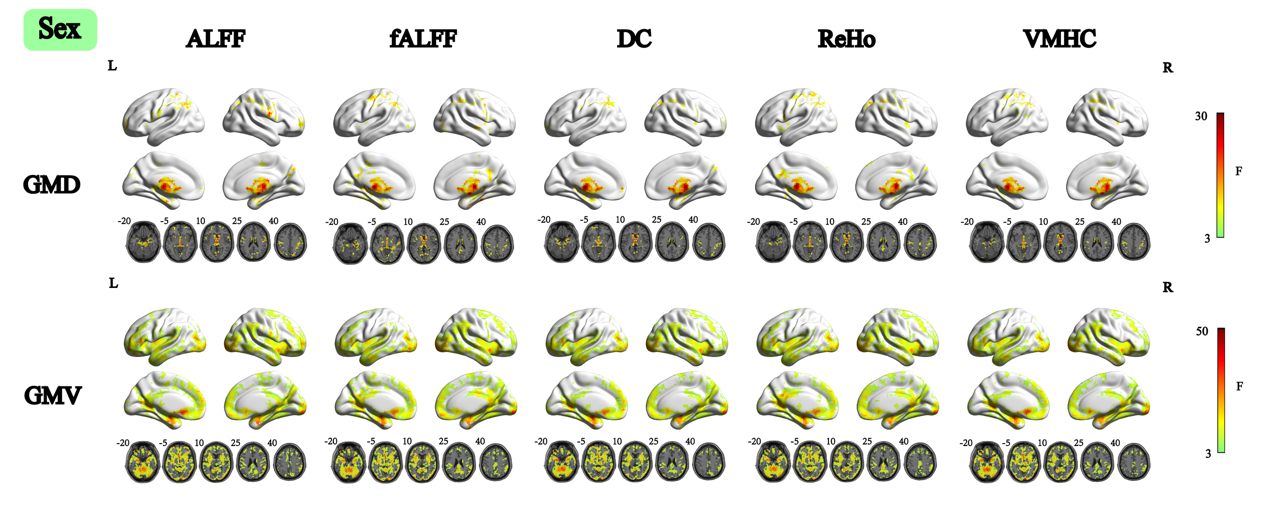

Fig. S2 F-test results of sex. GRF correction (voxel-level *p* < 0.001, cluster-level *p* < 0.05) was utilized to correct the large number of voxels across the brain.

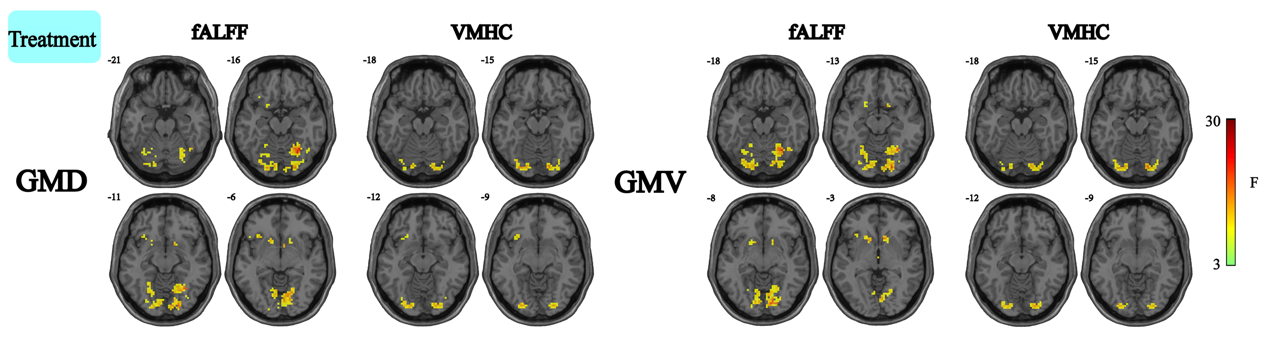

Fig. S3 F-test results of treatment time. GRF correction (voxel-level *p* < 0.001, cluster-level *p* < 0.05) was utilized to correct the large number of voxels across the brain.
